# Detection of Ovarian Cancer Using Samples Sourced from the Vaginal Microenvironment

**DOI:** 10.1101/597740

**Authors:** Melissa M. Galey, Alexandria N. Young, Valentina Z. Petukhova, Mingxun Wang, Jian Wang, Joanna E. Burdette, Laura M. Sanchez

## Abstract

Mass spectrometry (MS) offers high levels of specificity and sensitivity in clinical applications, and we have previously been able to demonstrate that matrix-assisted laser desorption/ionization time-of-flight (MALDI-TOF) MS is capable of distinguishing two-component biological mixtures at low limits of detection. Ovarian cancer is notoriously difficult to detect due to the lack of any screening methods for early detection. By sampling a local microenvironment, such as the vaginal fluids, a MS based method is presented that was capable of monitoring disease progression from vaginally collected, local samples from tumor bearing mice. A murine xenograft model of high grade serous ovarian carcinoma (HGSOC) was used for this study and vaginal lavages were obtained from mice on a weekly basis throughout disease progression and subjected to our MALDI-TOF MS workflow followed by statistical analyses. Proteins in the 4-20 kDa region of the mass spectrum could consistently be measured to yield a fingerprint that correlated with disease progression over time. These fingerprints were found to be statistically stable across all mice with the protein fingerprint converging towards the end point of the study. MALDI-TOF MS serves as a unique analytical technique for measuring a sampled vaginal microenvironment in a specific and sensitive manner for the detection of HGSOC in a murine model.

## Introduction

Ovarian cancer is a severe gynecological disease that is currently the fifth leading cause of cancer deaths among women in the United States, with an estimated 22,530 newly diagnosed cases and 13,980 deaths expected in 2019.^1^ This disease has a five-year survival rate of 93%, if diagnosed when tumor growth is limited to reproductive organs, such as the ovaries or fallopian tubes.^2,3^ However, the majority of diagnoses occur during later stages of the disease when tumors have already metastasized, resulting in a five-year survival rate of less than 30%. Around 90% of these diagnoses are of epithelial origin, of which high-grade serous ovarian cancer (HGSOC) is the most common subtype.^3–5^ These late-stage diagnoses can be attributed to the lack of any routine screening methods available for early detection of the disease. Current diagnostic methods, such as transvaginal ultrasounds and CA-125 biomarker tests, are generally used only after patients present with symptoms, and they are considered invasive procedures. These methods also suffer from inaccuracy and a lack of specificity, which limits their use in diagnosing ovarian cancer in its early stages.^6,7^

In a routine health screening, women receive a Pap smear to determine cervical health, but recent data suggests that these samples may be repurposed for endometrial and ovarian cancer detection.^8^ This, along with other studies, shows that DNA from ovarian cancer cells can be detected in clinical samples, such as Pap smears, which represents a concentrated sample from the local microenvironment in the female reproductive system.^8–10^ This clinical evidence would suggest that cells and/or cellular material from ovarian tumors reside outside of the proximal environment in areas such as the cervix or vagina fluid. Further confirmation of the Pap smear linked to HGSOC detection was recently detailed in a case report, in which adenocarcinoma cells derived from HGSOC precursor lesions were observed in a cervical smear.^11^ This case study indicates that developing a technology or measurement using Pap smears may allow for the detection of HGSOC.

While reliable early diagnostic markers for ovarian cancer have remained elusive, research has moved closer towards characterizing the proteome of a healthy female reproductive system using MS.^12^ For instance, the iKnife and MassSpec Pen are both innovative devices that couple MS signatures in lipids to the detection of cancerous tissue in a surgical setting.^13,14^ Work has recently been done using mass spectrometry (MS) to investigate Pap smears under the assumption that cells, cellular debris or secreted proteins from the female genital tract would be present.^12^ Bottom-up proteomics using a LTQ Orbitrap mass spectrometer was performed on liquid Pap smear samples from women considered to healthy to create a ‘Normal’ Pap Test Core Proteome.^12^ This analytical method is highly sensitive, but it requires significant sample preparation, such as sample cleanup, to remove salts and insoluble particulates, protein digestion, and lengthy liquid chromatography times leading to a time investment, as well as a high degree of expertise needed to operate a high mass resolution tandem mass spectrometer (LC-MS/MS). We chose to investigate the use of matrix-assisted laser desorption/ionization time-of-flight (MALDI-TOF) MS since this is considered a versatile analytical technique used in a variety of applications ranging from the characterization of microbial species to biomarker detection using protein signatures.^15–17^ For example, the MALDI-TOF Biotyper system is currently FDA approved for use in clinical diagnostic laboratories for microbial typing.^18^ MALDI-TOF MS has also been used to characterize mammalian cells based on spectral fingerprints, which in turn, allows for their identification.^19–21^ Perhaps in combination with genomic sequencing, the ability to directly detect proteins, particularly those that change in abundance over time from a local source, may improve strategies that can be leveraged for detection.

As previous studies have highlighted that ovarian cancer cells and associated cellular debris have the ability to travel through the female reproductive system to the cervix, we felt it would be feasible to detect these changes to the reproductive environment using MALDI-TOF MS based on whole-cell fingerprinting with protein signatures.^8,10,21^ As a proof of principle for this concept, we used a murine model of HGSOC. In this study, we used vaginal lavages (analogous to Pap smears) in a murine xenograft model to detect the increasing burden of ovarian cancer based on protein fingerprints in the 4-20 kDa mass range obtained using MALDI-TOF MS (**Figure 1**). Statistical analysis of the protein signatures found candidate proteins that both increase and decrease over time as disease progresses highlighting the importance of sampling a local microenvironment proximal to disease origin. By sourcing cells from a local microenvironment, we hoped to achieve greater sensitivity and specificity while also exploring the ability of small proteins to lend themselves as possible biomarkers of disease progression.

**Figure 1:**
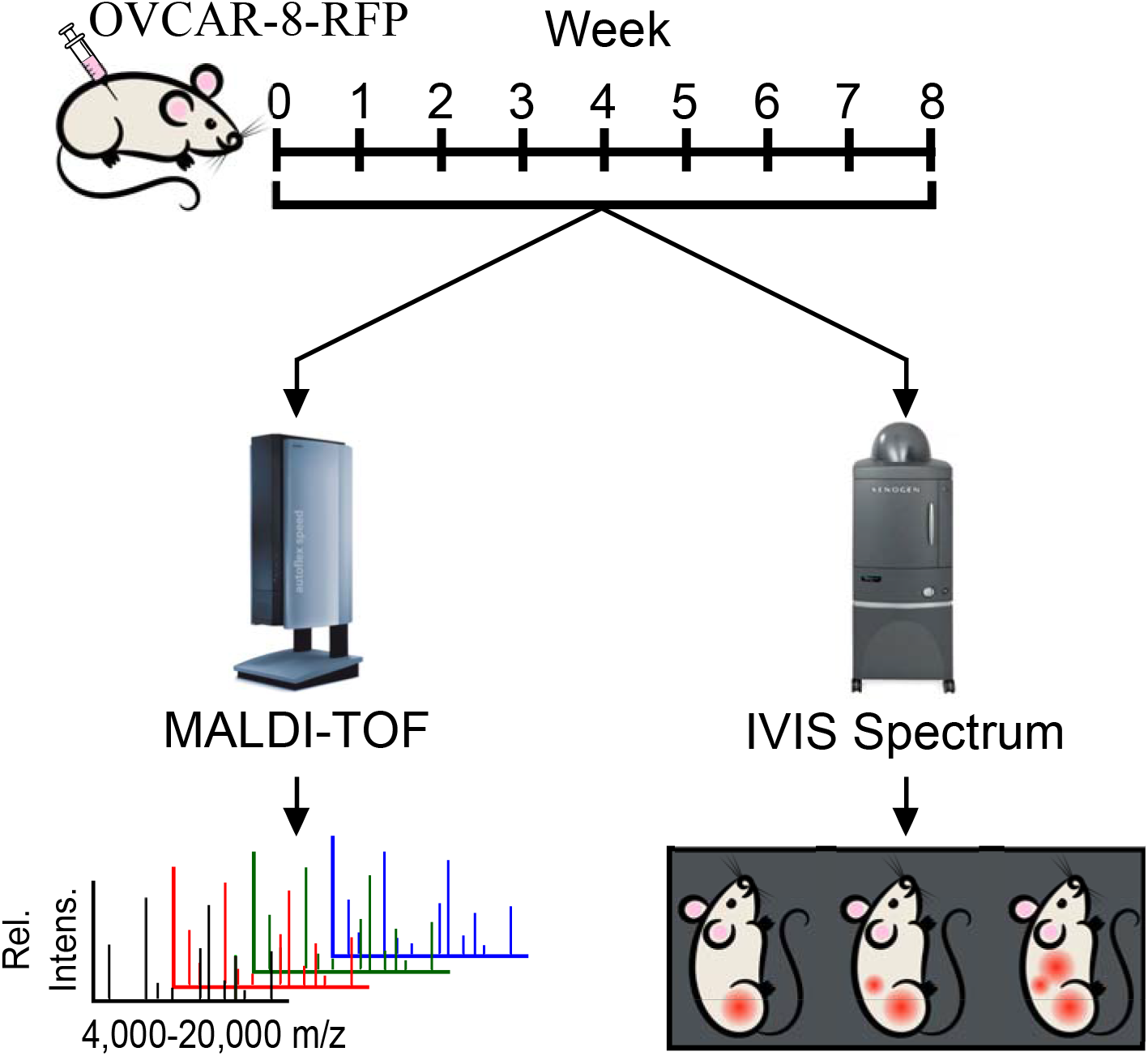
Workflow of murine xenograft study. Female athymic mice are IP injected with OVCAR-8-RFP and tumors were allowed to develop over an eight week period. Each week, mice were given vaginal lavages and were imaged using an IVIS imaging system to track tumor burden. Vaginal lavages were analyzed using MALDI-TOF MS to obtain mass spectra for statistical analyses.

## Materials and Methods

OVCAR-8, a human-derived ovarian cancer cell line, expressing red fluorescent protein was a gift from M. Sharon Stack at the University of Notre Dame and were STR validated at UIC. Additional information regarding the Materials and Methods section can be found in the **Supporting Information**.

### *In Vivo* Murine Xenograft Study

5×10^6^ OVCAR-8-RFP cells suspended in PBS were injected intraperitoneally into NCr *nu/nu* athymic female mice (N=5) aged 10 to 12 weeks (Taconic, Rensselaer, NY). Tumor burden was monitored on a weekly basis using a Xenogen IVIS^®^ Spectrum *In Vivo* Imaging System (PerkinElmer, Waltham, MA). Mice were also given weekly vaginal lavages using 200 μL of sterile PBS throughout the study to collect cells from the local microenvironment of the reproductive organs. Cells sourced from lavages were counted, normalized to 10,000 cells/μL using deionized water and stored at −80°C following collection. After two months of tumor progression, all animals were humanely sacrificed followed by collection of tumors and reproductive organs.

### MALDI-TOF MS

Frozen lavage suspensions were thawed on ice and diluted to 5,000 cells/μL using deionized water. Equal volumes of lavage samples and a 20 mg/mL sinapic acid matrix solution (Sigma Aldrich, St. Louis, MO) in 30/70 ACN/H_2_O with 1% TFA were mixed together for a final concentration of 2,500 cells/μL. Samples were placed on ice for ten minutes and 1.5 μL of each sample was spotted onto a 384- well ground steel MALDI target plate with 24 technical replicates per lavage with Protein Standard I (Bruker Daltonics, Billerica, MA) used as a calibrant. This procedure was previously optimized and described in depth in Petukhova *et. al.*^21^ An Autoflex Speed LRF MALDI-TOF mass spectrometer (Bruker Daltonics, Billerica, MA) was used to acquire mass spectra of murine vaginal lavages samples in positive linear mode with a mass range of 4 to 20 kDa using a laser power of 75%, a laser width of 3 (medium) and a gain of 18.1x. Protein spectra was acquired using AutoXecute, an automated run function of FlexControl v. 3.4 software (Bruker Daltonics, Billerica, MA). 4,000 laser shots were accumulated in fifty shot increments for each sample using the random walk option. All mass spectra were externally calibrated using Protein Standard I (Bruker Daltonics, Billerica, MA), which had been spotted directly adjacent to each sample spot in a square pattern.

Spectra were pre-processed using FlexAnalysis v. 3.4 (Bruker Daltonics, Billerica, MA) with baseline subtraction (TopHat algorithm) and smoothing (SavitzkyGolay algorithm, width of *m/z* 5 over five cycles). Peak picking was performed using a signal-to-noise threshold of four following processing. Mass spectra were converted to a mzML format and further pre-processed using MALDIquant, an open-source R package, (https://cran.r-project.org/web/packages/MALDIquant/MALDIquant.pdf) using a standard workflow consisting of steps such as baseline correction and spectra alignment; resulting in a feature matrix comprised of intensity and *m/z* values shown in (**Figure S1**). Spectral data are publicly available online (ftp://MSV000083628@massive.ucsd.edu).

### Statistical Analysis

In order to search for features that were differentially expressed throughout the longitudinal sampling points in every mouse, a Wilcoxon rank-sum test was applied, using the abundances of each feature, to assess whether the abundance differences between all pairs of days within one mouse were statistically significant. The Wilcoxon rank-sum test was chosen because it makes very few assumptions about the distribution of the data, e.g. normality in the case of t-tests. A Bonferroni-correction was applied to the p-value of each test to account for multiple hypothesis testing and a threshold of 0.01 was applied on the corrected p-value for each test. Source code is available online (https://github.com/mwang87/ometa_mousemaldi).

## Results and Discussion

### *In Vivo* Imaging to Monitor Tumor Progression in OVCAR-8 Xenograft Model

HGSOC was modeled *in vivo* using an orthotopic xenograft model by intraperitoneal injection of the OVCAR-8-RFP human ovarian carcinoma cell line. OVCAR-8 harbors a p53 mutation, and p53 mutation is reported in up to 96% of HGSCO tumors.^22–24^ Following the IP injection of OVCAR-8-RFP cells, all mice were imaged every seven days and their respective fluorescence increased in intensity and colonized the peritoneum over time (**Figure 2** **and S2**). At the conclusion of the longitudinal study, all five of the mice had significant metastatic tumor burden and four out of the five presented with ascites, a buildup of fluid in the abdominal cavity, which is a symptom present in over a third of ovarian cancer patients.^25^

**Figure 2:**
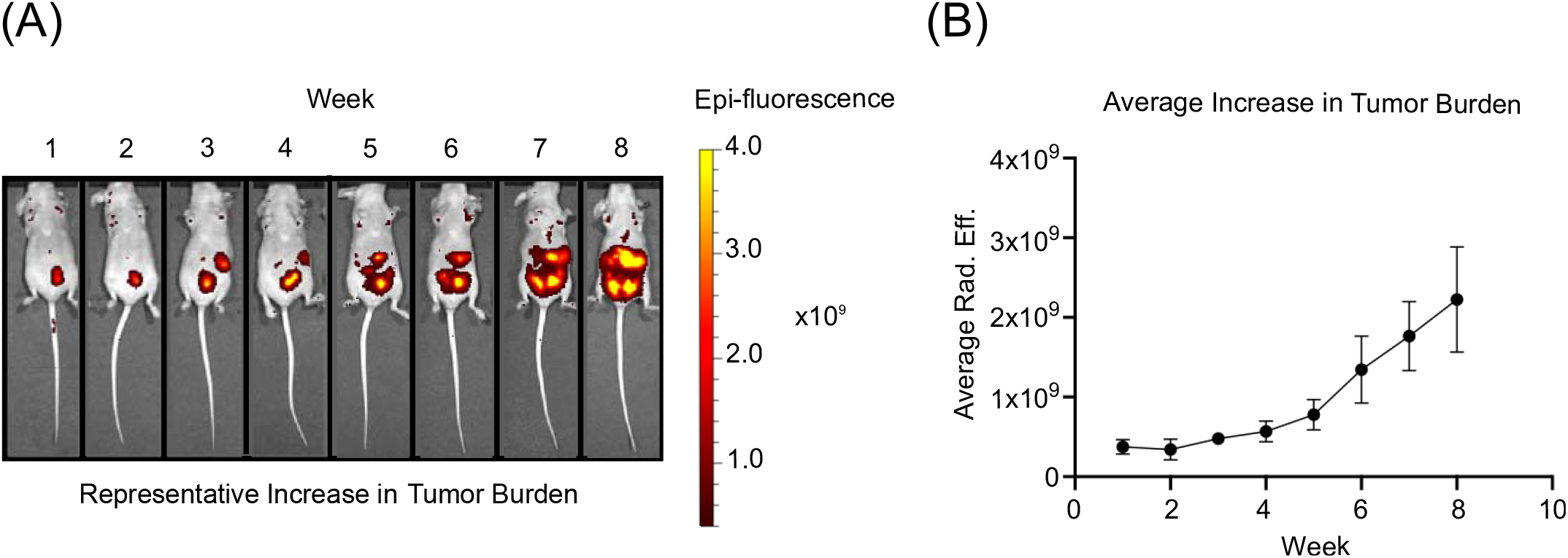
(A) *In vivo* fluorescence imaging of OVCAR-8-RFP tumors in athymic nude mice over an eight week period (B) Average radiant efficiency over time for all mice indicating an increase in tumor burden.

### MALDI-TOF MS Offers a Lower Limit of Detection than Fluorescence Imaging

Previous work in our lab by Petukhova *et. al* has demonstrated that whole-cell fingerprinting, using MALDI-TOF MS, is capable of detecting signals from 1% of lysed cancer cells in an otherwise healthy heterogenous two-component mixture.^21^ Spiking studies performed using lysates of healthy murine vaginal lavages and freshly harvested OVCAR-8-RFP cells indicated that an otherwise healthy cell mixture would need to be comprised of at least 10% OVCAR-8-RFP cells in order to reproducibly detect fluorescence (**Figure S3**). This would indicate that MALDI-TOF MS is a more sensitive and reproducible technique for detecting changes in the local microenvironment where the biological sample was sourced, such as the reproductive proteome from a murine vaginal lavage. This also suggests that detection of cellular components, rather than intact cells, is feasible.

We had originally anticipated that the fluorescent tag on the OVCAR-8-RFP cells would allow for cell counting in the vaginal samples throughout the study. However, based on the outcome of the spiking study, it would be improbable to see 10% of the cells in the samples as the RFP tagged ones during a routine vaginal lavage and unlikely that fluorescence imaging would be of use for detecting these types of cells in our samples. While we did not observe fluorescence in murine lavages at any time point, changes were detected in the protein fingerprint over time using MALDI-TOF MS, which is of interest as changes in the local microenvironment were occurring as the disease progresses. As cells were lysed prior to analysis, our MS method relies on the detection of proteins from lavages instead of whole cells, making the source of these proteins ambiguous. For instance, we can not differentiate between cellular origination from tumor cells versus those found in the murine reproductive system. Moreover, it could also be possible that we were observing the cellular response to cancer, but this provided a unique fingerprint as compared to mice with another peritoneal disease. The observance of statistically significant changes in feature intensity, regardless of whether the proteins were from ovarian cancer tumor cells or not, is of importance for this study as we were not focused on the appearance of tumor cells in the murine lavage, but rather changes in the overall system and/or cellular components.

### Vaginal Lavages from a HGSOC Model Yield Distinct Differences from Other Disease Models

Based on our previous work with two-component mixtures with different cell lines, it was anticipated that a vaginal lavage taken over time would be useful for monitoring ovarian cancer progression.^21^ In order to verify that these protein fingerprints are unique to cancer and not a result of peritoneal disease, mice with non-alcoholic steatohepatitis (NASH), which results in severe liver damage, were age-matched to those in our OVCAR-8 xenograft model in its seventh week and vaginal lavages were collected. Upon comparison of average spectra for each condition, it was observed that the protein signatures were distinct between the two peritoneal diseases (**Figure S4**). Therefore, our MALDI-TOF MS methodology yields distinct, disease specific protein fingerprints from the vaginal microenvironment. Additionally, while our study consisted of vaginal lavages on a weekly basis, we wanted to ensure that significant changes to the protein fingerprint are truly due to the progression of HGSOC and not inflammation of the vaginal cavity. Repetitive vaginal lavages were performed on healthy athymic nude mice over a five day period and protein signatures were not significantly different from day-to-day (**Figure S5**). It is also important to note that the murine estrous cycle spans a period of four to five days so slight changes could be attributed to this phenomenon, but the cycle was not sufficeint to drive alterations in the fingerprint.

### Use of MALDI-TOF for Analysis of Vaginal Lavage Samples

MALDI-TOF MS had several advantages for this study such as solvent-free, rapid data acquisition (milliseconds), minimal sample requirement, and the availability of this technology in clinical laboratories.^15–18^ It also allows rapid collection of a protein fingerprint over a large mass range, providing intact protein masses. Similar to how the Biotyper system (Bruker Daltonics) works for microbial identifications, the collection of a protein fingerprint allows for the comparison of different spectral patterns, in terms of the appearance and absence of features, to look for matches based on characteristic peaks.^18^ This is advantageous to our study as we have the ability to statistically analyze and compare protein fingerprints collected at various time points throughout the study to look for changes in protein peaks during tumor progression. Our lab had previously identified the small protein mass range (4-20 kDa) to a higher number of characteristic peaks as well as the most distinctive when comparing cell types, particularly human ovarian cancer cell lines and murine oviductal cell lines.^21^ Therefore, protein fingerprints from this mass range were collected for murine vaginal lavages from weeks one through seven of the xenograft study, as they were most likely to yield the richest spectral data.

In order to begin to visualize the data, the cosine similarity of the murine vaginal lavages in the form of a dot product matrix was used (**Figure 3**). This allows for an easily accessible view of how the profiles may differ over time when compared to specific weeks in the study. At the beginning of the study, each biological replicate (N=5) had a largely unique profile when compared to others at or around the same time point (red dots) with low similarity being observed. However, one of the striking results was that at the the end of the study during the seventh week, the protein fingerprints from the vaginal lavages are largely conserved (green dots), which would indicate that all five mice have similar protein profiles once the tumor burden is representative of late stage ovarian cancer. This would imply that there are unique processes and/or cellular responses associated with the progression of HGSOC that can be captured in a spectral protein fingerprint. All detected peaks that were found to be differentially expressed across time points can be found in the Supporting Information (**Table S1**).

**Figure 3:**
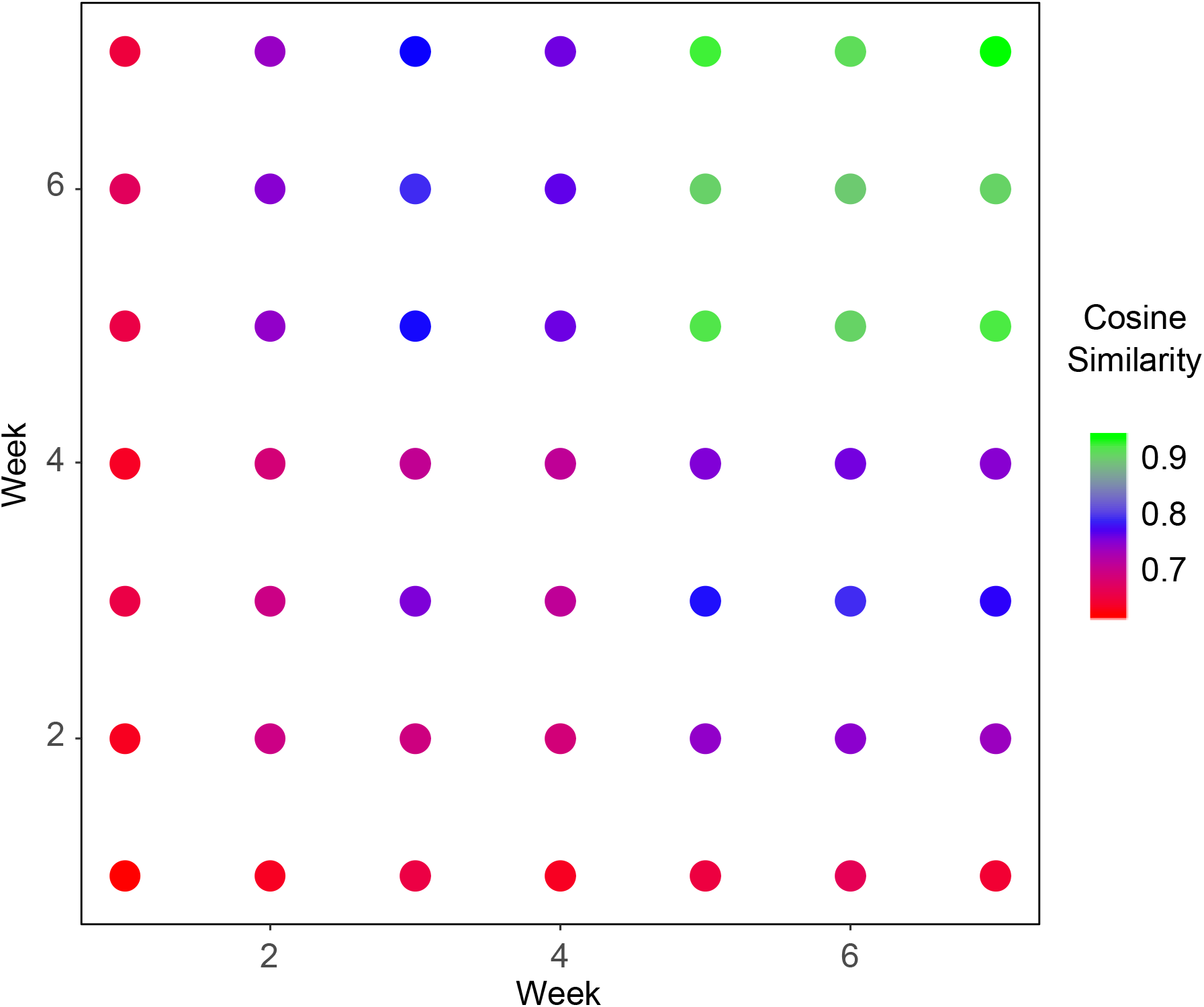
Dot matrix plot comparing spectral similarity between time points. High cosine scores (green) indicates that the global profiles are highly similar between intersecting time point (i.e. weeks six and seven) while low cosine scores (red) indicates less similarity in features between the spectra of the respective time points.

This late stage similarity prompted us to search for the presence and absence of protein features across all mice across paired time points (i.e. week one versus week seven). We found that multiple proteins began to dominate the mass spectra throughout disease progression and that a larger number of proteins either appeared and become absent from the protein fingerprint over time. This led to the investigation of specific regions of the spectrum and the verification that specific peaks became either significantly up- or downregulated over time, which would suggest that tumor progression does, in fact, induce change in specific processes in our murine model system. Several research groups have studied the proteome of cervical-vaginal fluid obtained from women found to be in good health and while some of the proteins they were identified fell within the mass range detailed in this study, such as thioredoxin and profilin-1, there is not much known about the reproductive proteome between *m/z* 4,000-20,000, prompting additional research.^12,26,27^ Some have speculated that this mass range contains an abundance of histones or ribosomal proteins based on previous whole-cell fingerprinting research and we hope to expand the knowledge of this area in the future through the identification of detected proteins using MS.^19^

The Wilcoxon-rank sum test was used for the normalized intensity values from each observed peak and ranks them in terms of value for comparison. This statistical test makes minimal assumptions about the data, which allows it to be used in cases of non-normal distributions. Following analysis, specific features, shared between all mice, were found to be significantly change in intensity throughout the time course of the study (**Figure 4**). The determination of the test statistic in each comparison (week to week) allowed for the calculation of a p-value for significance. This analysis allowed for the identification of 118 upregulated and 97 downregulated spectral features when comparing data from different time points, implying that there were multiple significant changes over time in our mouse model. In particular, there are 19 *m/z* values and 10 *m/z* values that are significantly up- and downregulated, respectively, in all mice (p <0.001). Interestingly, the significant features between the initial and final time points of the study are segregated into different subsets of the mass range (**Figure S6**). The majority of the signals that appear over time are approximately twice the mass as those observed to be downregulated. This may point to a dysregulation in protease activity and requires further validation on the identity of these proteins to further test the possible mechanism by which these proteins accumulate in the disease state.

**Figure 4:**
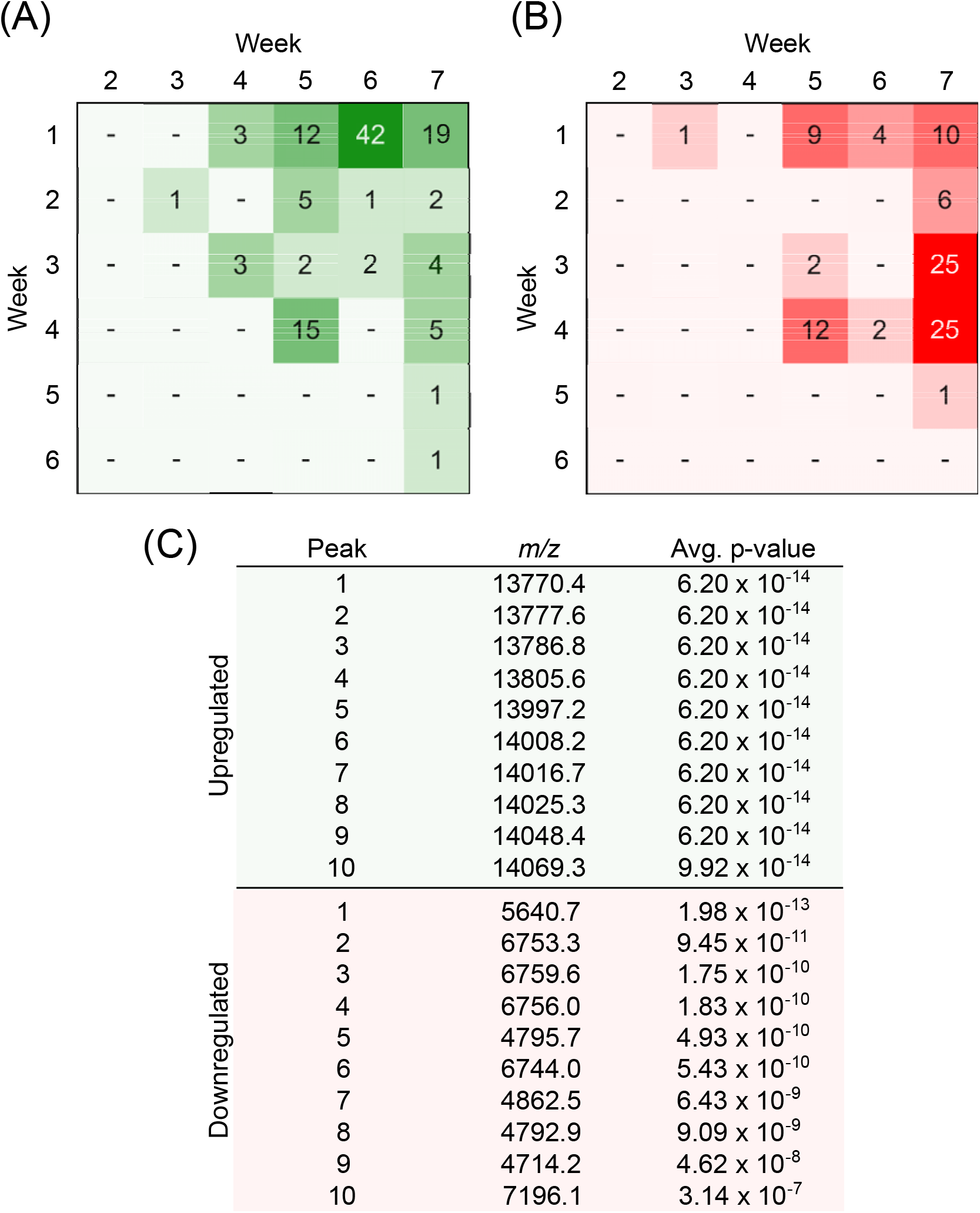
(A) Pivot table showing number of significantly upregulated signals observed across time points across all five mice. (B) Pivot table showing number of significantly downregulated signals observed across time points across all five mice. (C) Top ten *m/z* values according to average p-values when comparing weeks one and seven.

While the spectral fingerprint obtained from the murine vaginal lavages is a promising start to better understanding the cellular processes that occur throughout the progression of ovarian cancer, it is of note that while MALDI-TOF mass spectrometers allow for the observance of intact protein masses, they do not have the mass resolving power in linear mode to provide an exact mass for the detected spectral features. Bottom-up proteomics, performed on a high-resolution instruments such as a hybrid quadrupole-orbitrap, will be required to definitively identify these proteins via analysis of peptide fragments. Bottom-up proteomics has its own limitations such as potentially not having enough cleavage sites for the digestions enzyme due to the small size of the proteins. However, using a combination of both MS methods to obtain an intact mass along with tandem fragmentation will allow us to identify these proteins in the future.

## Conclusion

Based on statistical analysis with the Wilcoxon rank-sum test, we identified intact *m/z* values of small proteins obtained from murine vaginal lavages. We conclude that the protein fingerprint region obtained from our MALDI-TOF MS technique can be used to correlate with disease progression of ovarian cancer *in vivo*. By sourcing samples from a local microenvironment, such as the vaginal cavity or cervix, we have shown that significant changes in protein signatures can be observed over time in the progression of ovarian cancer in a murine model. Our next step will be to further investigate and identify these proteins of interest using tandem MS to gain a better understanding of their role in disease progression in our *in vivo* model. A deeper understanding of these signature proteins and the processes they are involved in may further enhance our ability to design a diagnostic test.

## Author Contributions

JEB and LMS were responsible for the conception of the study. MMG and ANY were responsible for cell and animal work, including vaginal lavages, imaging and tumor collection following sacrifice. MMG and VZP were responsible for MS data collection and analysis. MW and JW were responsible for performing the statistical analysis required for this study. All authors were responsible for preparation of the manuscript with input and review received from all authors.

## Notes

Dr. Mingxun Wang is a consultant for Prometheus Laboratories Inc., a diagnostics company. However, Dr. Wang’s involvement with Prometheus Laboratories Inc. is not related to the material in the submitted work.

## Acknowledgements

This research was supported by the National Center for Advancing Translational Sciences, National Institute of Health, through grant UL1TR002003 (JEB & LMS), the Research Corporation for Science Advancement Scialog® award #26222 (LMS),University of Illinois at Chicago Start-up Funds (LMS), UG3 ES029073 (JEB), and F30 CA217079 (ANY). The authors would also like to acknowledge Daniel D. Lantvit, a research specialist in the Burdette lab, for his expertise and assistance with initiating our *in vivo* work. We would also like to acknowledge Dr. Natalia Nieto at UIC for allowing us to collect samples during her animal study concerning NASH.

