## Supporting Information for "Detection of Ovarian Cancer Using Samples Sourced from the Vaginal Microenvironment"

| **Table of Contents** | |
| --- | --- |
| Supporting Information: Materials & Methods…………………………………………................ | S2 |
| Figure S1: Pre-processing Workflow Using the MALDIquant Package in R..………………… | S3 |
| Figure S2: *In vivo* Fluorescence Imaging of OVCAR-8-RFP Tumors in Athymic Mice………. | S4 |
| Figure S3: Microscopy Images of RFP Spiking Study…………………………………………… | S5 |
| Figure S4: Mirror Plot Comparing Protein Fingerprints from Week Seven in a HGSOC and NASH Model………………………………………………………………………………..………… | S6 |
| Figure S5: Mass Spectra of Repeat Murine Lavages from Healthy Mice …………………..… | S7 |
| Figure S6: Mirror Plot Comparing Protein Fingerprints from Weeks One and Seven in a HGSOC Model……………………………………………………………………………………….. | S8 |
| Table S1: Up- and Downregulated Spectral Features Differentially Expressed Across Various Time Points Throughout the Study……………………………………………………….. | S9 |

**Supporting Information: Materials & Methods**

**Cell Culture**

OVCAR-8-RFP were grown in DMEM with 10% FBS and 1% penicillin/streptomycin. Cultured cells were maintained in a humidified incubator at 37°C in 5% CO_2_. Cells were passaged a maximum of 20 times. Cell lines were validated by short tandem repeat analysis and tested mycoplasma-free in 2017.

***In Vivo* Murine Xenograft Study**

Tumor burden was monitored using a Xenogen IVIS^®^ Spectrum *In Vivo* Imaging System (PerkinElmer) as described by Lewellen *et al.* Imaging was done using an exposure time of one second, an Fstop of two, and an excitation and emission wavelength of 535 and 620 nm, respectively.^26^ Vaginal lavage samples using 200 μL of sterile PBS as described by McLean *et. al*. at various time points prior to xenografts to collect healthy samples.^25^ Mice were housed in facilities managed by the Biological Resources Laboratory at UIC and were provided food and water *ad libitum*. All animals were humanely treated in accordance with the Animal Care and Use Committee guidelines at the University of Illinois at Chicago (UIC) using Protocol #17-174.

Vaginal lavage samples were also collected using 200 μL of sterile PBS of Black 6 age-matched mice (N=9) with NASH for comparative purposes to our ovarian cancer study.

**Cell Counting for MS Analysis**

Cells sourced from murine vaginal lavages were counted using a K2 Cellometer (Nexcelom, Lawrence, MA) by pipetting 20 μL of the sample into disposable counting chambers for imaging. Brightfield images were taken of each sample for accurate counts of leukocytes and epithelial cornified cells present which were used to calculate the concentration of the lavage samples using FCS Express software (De Novo). Fluorescent images were also taken using a 660 nm filter to determine if OVCAR-8-RFP cells were present in murine samples. All lavages were spun down at 150 rcf for five minutes, at which point, excess PBS was removed. Cells were resuspended in deionized water to reach a final concentration of 10,000 cells/μL.

**Limit of Detection Spiking Studies**

OVCAR-8-RFP cells as well as cells sourced from murine vaginal lavages were separately counted using a K2 Cellometer (Nexcelom, Lawrence, MA) and concentrated to a final concentration of 10,000 cells/μL in deionized water. RFP-tagged cells were spiked into healthy cell mixtures at 1% and 10% for detection. Fluorescent images were taken using a 660 nm filter to quantify the number of OVCAR-8-RFP cells present in the lavage samples.

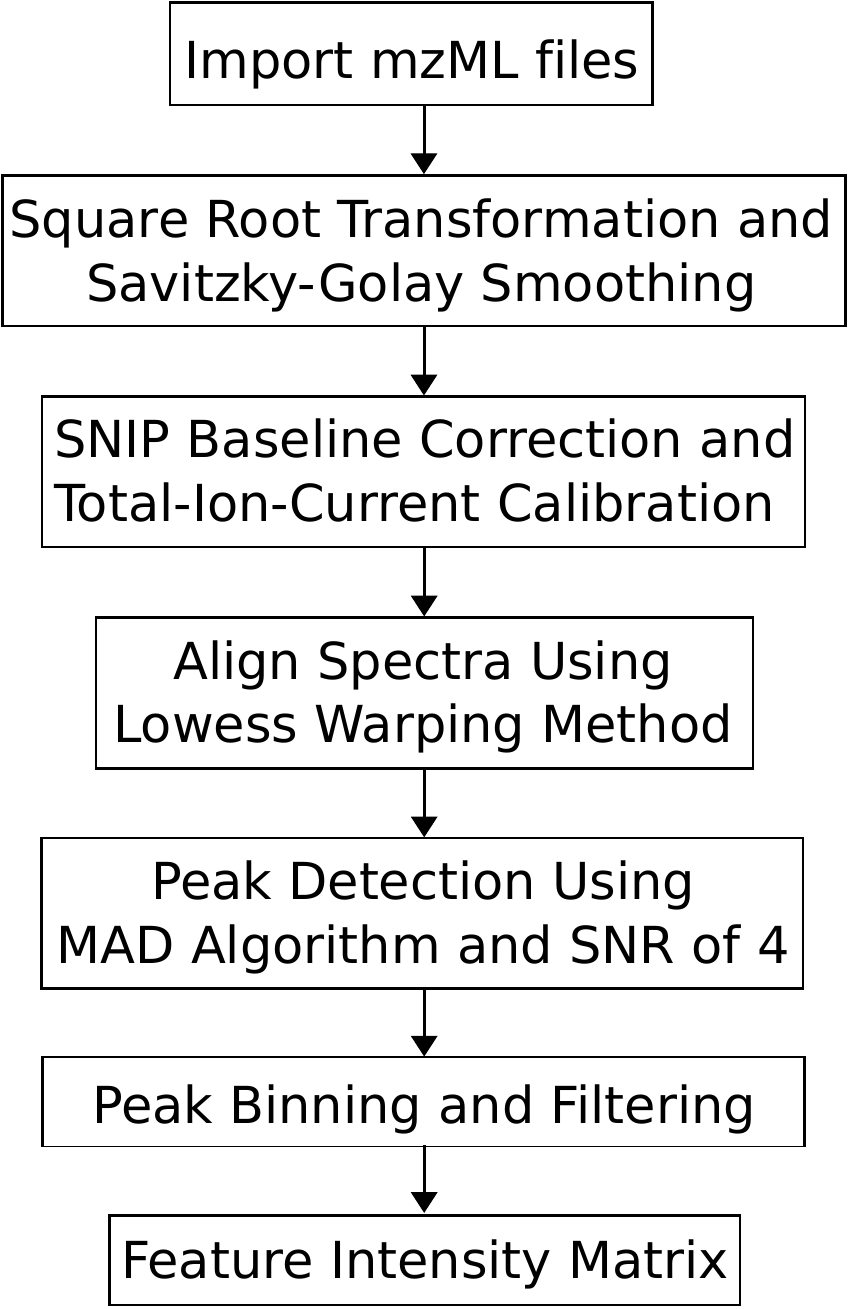

Figure S1: Pre-processing workflow using the MALDIquant package in R. All mzML files were uploaded using the MALDIquant package and batch processed using the outlined parameters, resulting in a feature matrix that consists of feature peaks and their corresponding intensity values.

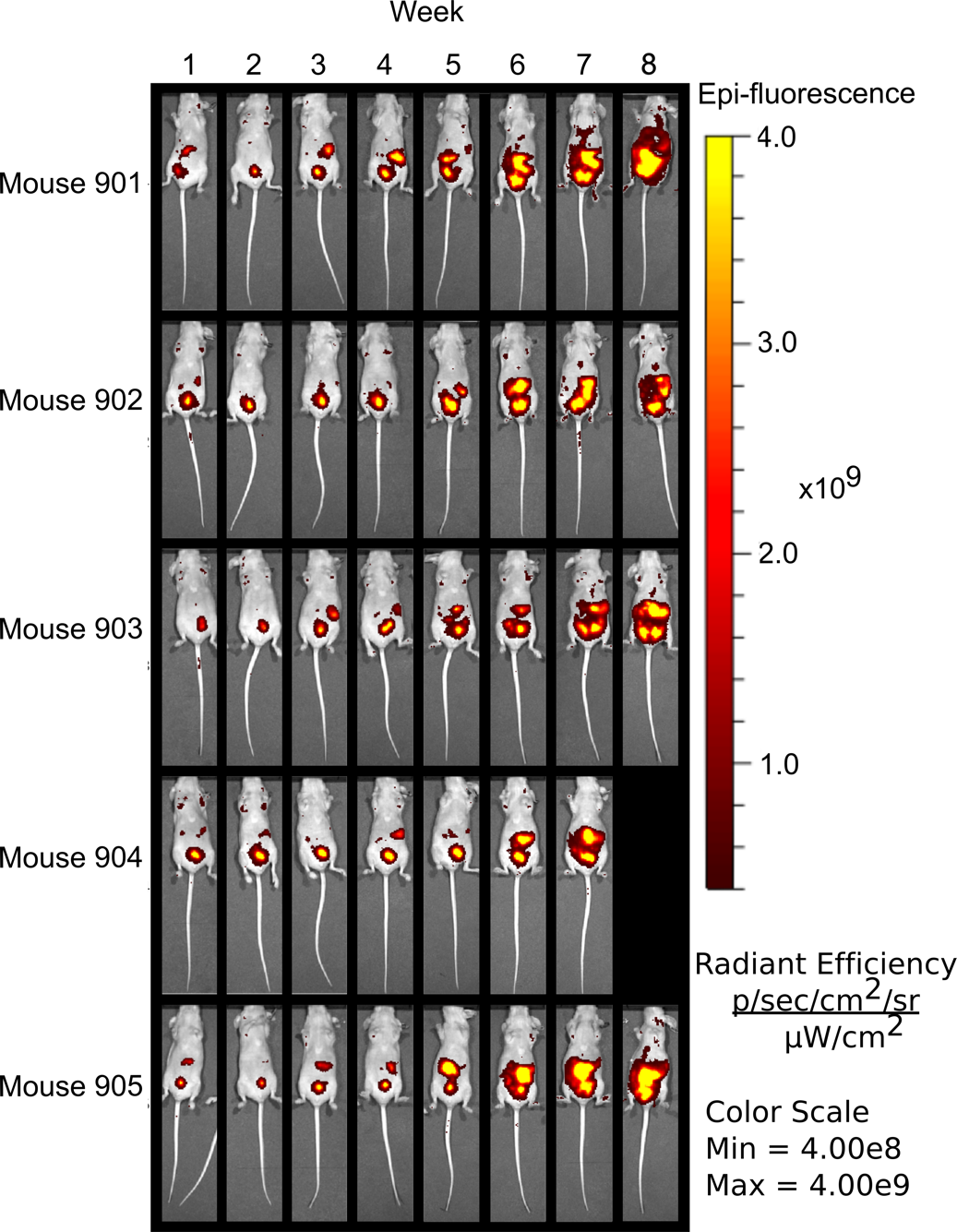

Figure S2: *In vivo* fluorescence imaging of athymic nude mice across the time course of the study showing OVCAR-8-RFP tumor progression. Mice were imaged on a weekly basis for a time span of two months. Mouse 904 expired prior to the conclusion of the study, which is why no image was obtained during the eighth week.

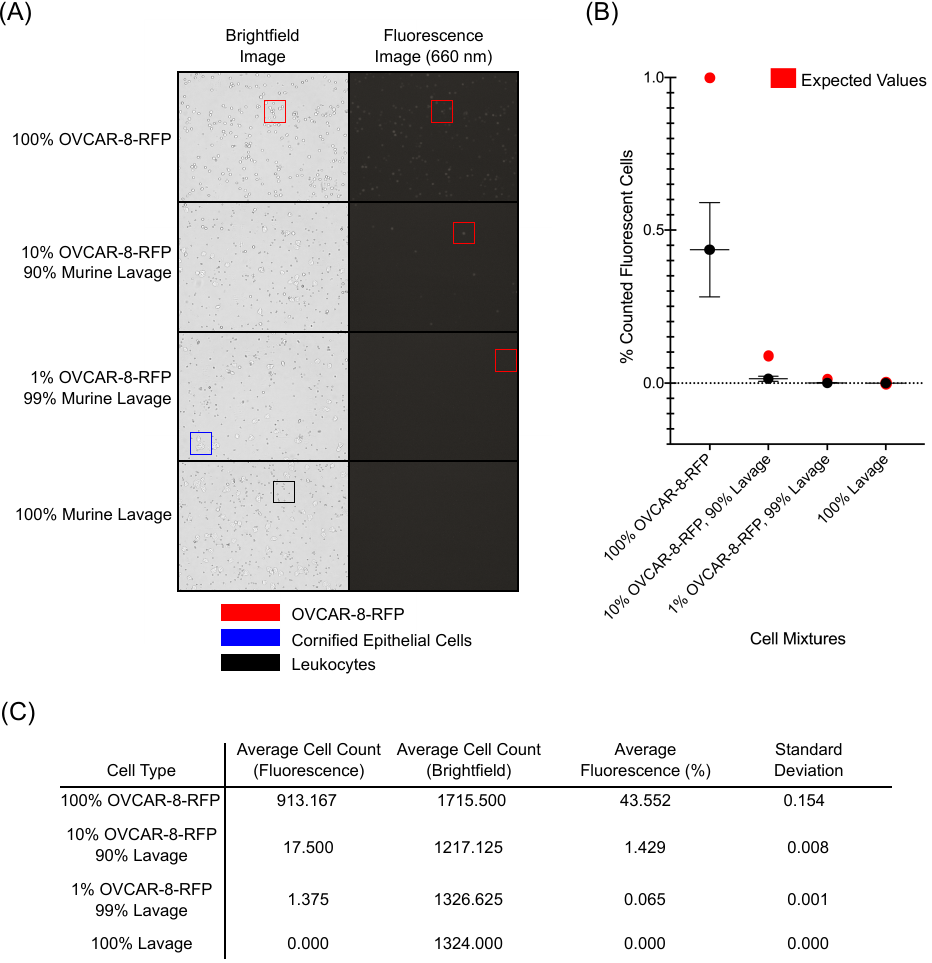

Figure S3: (A) Representative brightfield and fluorescence (600 nm) images of OVCAR-8-RFP cells, murine lavages and mixtures of the two components with red boxes outlining examples of fluorescent cells. (B) Overall fluorescent cell count as a percentage of cells detected from brightfield images via K2 Cellometer (Nexcelom, Lawrence, MA) compared to the expected values. (C) Table detailing the average cell count (fluorescent and brightfield) along with average fluorescence and respective standard deviations (n=6).

Figure S4: Mirror plot of average protein fingerprints of murine lavages comparing week seven in an ovarian cancer model and week seven in a NASH model (N=5, n=24 and N=9, n=24, respectively). (A) Full spectra (*m/z* 4,000-20,000) with blue and green sections indicating regions of interest. (B) Spectral features in this region (*m/z* 4,600-5,400) differ between both disease states in terms of peaks and intensity. (C) Spectral features in this region (*m/z* 10,000-11,500) present another area in which peaks differ between both disease states in terms of appearance and intensity.

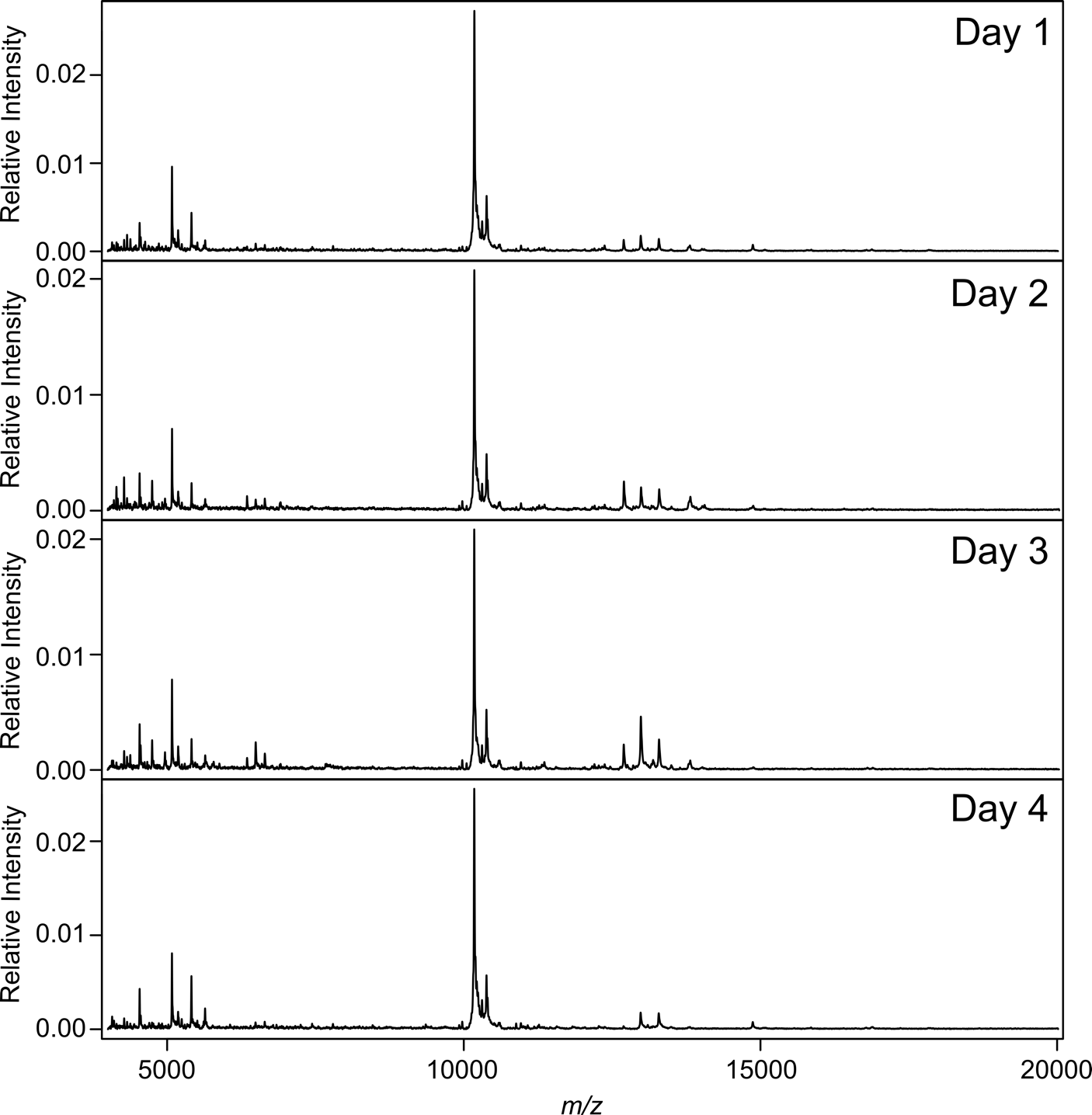

Figure S5: Vaginal lavages from healthy athymic nude mice were collected for a week on a daily basis. This was done to ensure that the signature detected at the conclusion of the study, when mice had heavy HGSOC tumor burden, was not due to inflammation of the vaginal cavity caused by repetitive sampling. It is also of note that the estrous cycle for mice lasts four to five days as that is the time period covered in this experiment.

Figure S6: Mirror plot of average protein fingerprints of murine lavages comparing weeks one and seven (N=5, n=24). (A) Full spectra (*m/z* 4,000-20,000) with blue and green sections indicating regions of interest. (B) Spectral features in this region (*m/z* 4,500-7,000) are upregulated between the two time points. (C) Spectral features in this region (*m/z* 12,500-14,500) are downregulated between the two time points.

Table S1: Pivot tables indicating spectral features that are differentially expressed across various time points throughout the study. (A) Upregulated *m/z* values with a p-value < 0.001 comparing weeks one through seven.  (B) Downregulated *m/z* values with a p-value < 0.001 comparing weeks one through seven.

| (A) Upregulated Features Shared Across all Five Mice | | | | | |
| --- | --- | --- | --- | --- | --- |
|  | **Week** | **3** | **5** | **6** | **7** |
|  |  | Unique Features | | | |
| **Week 1** | *m/z* | 1 | 9 | 4 | 10 |
|  | 4265.9 | - | - | 1 | - |
|  | 4269.8 | - | 1 | - | - |
|  | 4270.5 | - | 1 | - | - |
|  | 4367.7 | - | 1 | - | - |
|  | 4369.6 | - | 1 | - | - |
|  | 4714.1 | - | 1 | - | 1 |
|  | 4792.9 | - | - | - | 1 |
|  | 4795.7 | - | - | - | 1 |
|  | 4862.5 | - | - | - | 1 |
|  | 5508.8 | - | 1 | 1 | - |
|  | 5640.7 | - | 1 | - | 1 |
|  | 5807.5 | - | - | 1 | - |
|  | 5843.8 | - | - | 1 | - |
|  | 6744.0 | - | - | - | 1 |
|  | 6753.4 | - | - | - | 1 |
|  | 6756.0 | - | 1 | - | 1 |
|  | 6759.6 | - | - | - | 1 |
|  | 7196.1 | - | - | - | 1 |
|  | 12620.2 | 1 | 1 | - | - |
| **Week 2** | *m/z* | 0 | 0 | 0 | 6 |
|  | 6634.8 | - | - | - | 1 |
|  | 6661.3 | - | - | - | 1 |
|  | 6744.0 | - | - | - | 1 |
|  | 6749.6 | - | - | - | 1 |
|  | 6753.4 | - | - | - | 1 |
|  | 6759.6 | - | - | - | 1 |
| **Week 3** | *m/z* | 0 | 2 | 0 | 25 |
|  | 4048.1 | - | - | - | 1 |
|  | 4350.4 | - | - | - | 1 |
|  | 4358.6 | - | - | - | 1 |
|  | 4381.7 | - | - | - | 1 |
|  | 4567.6 | - | - | - | 1 |
|  | 4586.3 | - | - | - | 1 |
|  | 4586.8 | - | - | - | 1 |
|  | 4587.9 | - | - | - | 1 |
|  | 4612.7 | - | - | - | 1 |
|  | 4617.0 | - | 1 | - | 1 |
|  | 4631.0 | - | - | - | 1 |
|  | 4746.9 | - | - | - | 1 |
|  | 4753.1 | - | - | - | 1 |
|  | 4757.0 | - | - | - | 1 |
|  | 4759.0 | - | - | - | 1 |
|  | 4762.7 | - | - | - | 1 |
|  | 4952.0 | - | - | - | 1 |
|  | 5405.3 | - | - | - | 1 |
|  | 5412.6 | - | - | - | 1 |
|  | 5640.7 | - | - | - | 1 |
|  | 5645.6 | - | - | - | 1 |
|  | 6744.0 | - | - | - | 1 |
|  | 6756.0 | - | - | - | 1 |
|  | 6759.6 | - | - | - | 1 |
|  | 6767.4 | - | - | - | 1 |
|  | 9370.3 | - | 1 | - | - |
| **Week 4** | *m/z* | 0 | 12 | 2 | 25 |
|  | 4139.5 | - | 1 | - | - |
|  | 4140.7 | - | 1 | 1 | 1 |
|  | 4198.4 | - | - | - | 1 |
|  | 4226.2 | - | - | - | 1 |
|  | 4230.0 | - | - | - | 1 |
|  | 4269.8 | - | 1 | - | - |
|  | 4358.6 | - | 1 | - | - |
|  | 4381.7 | - | 1 | - | - |
|  | 4545.4 | - | 1 | - | - |
|  | 4566.7 | - | - | - | 1 |
|  | 4567.6 | - | 1 | - | 1 |
|  | 4617.0 | - | - | - | 1 |
|  | 4627.5 | - | - | - | 1 |
|  | 4631.0 | - | - | - | 1 |
|  | 4636.2 | - | - | - | 1 |
|  | 4757.0 | - | 1 | - | 1 |
|  | 4834.7 | - | 1 | - | - |
|  | 5412.6 | - | - | - | 1 |
|  | 5528.8 | - | 1 | - | - |
|  | 5640.7 | - | - | - | 1 |
|  | 6041.4 | - | 1 | - | 1 |
|  | 6233.9 | - | - | - | 1 |
|  | 6236.0 | - | - | - | 1 |
|  | 6653.0 | - | - | - | 1 |
|  | 6661.3 | - | - | - | 1 |
|  | 6759.6 | - | - | - | 1 |
|  | 6767.4 | - | - | - | 1 |
|  | 6838.4 | - | 1 | - | - |
|  | 8261.0 | - | - | 1 | - |
|  | 9354.6 | - | - | - | 1 |
|  | 18704.5 | - | - | - | 1 |
|  | 18716.8 | - | - | - | 1 |
|  | 18720.6 | - | - | - | 1 |
|  | 18767.5 | - | - | - | 1 |
| **Week 5** | *m/z* | 0 | 0 | 0 | 1 |
|  | 9947.8 | - | - | - | 1 |

| (B) Downregulated Features Shared Across all Five Mice | | | | | | |
| --- | --- | --- | --- | --- | --- | --- |
|  | **Week** | **3** | **4** | **5** | **6** | **7** |
|  |  | Unique Features | | | | |
| **Week 1** | *m/z* | 0 | 3 | 12 | 42 | 19 |
|  | 5110.5 | - | - | - | 1 | - |
|  | 5127.1 | - | - | - | 1 | - |
|  | 5129.8 | - | - | - | 1 | - |
|  | 5147.2 | - | - | - | 1 | - |
|  | 5149.1 | - | - | - | 1 | - |
|  | 6574.9 | - | - | - | - | 1 |
|  | 6904.0 | - | - | 1 | 1 | 1 |
|  | 7024.0 | - | - | 1 | - | 1 |
|  | 8179.0 | - | 1 | - | - | - |
|  | 10105.1 | - | - | - | 1 | - |
|  | 10108.1 | - | - | - | 1 | - |
|  | 10111.1 | - | - | - | 1 | - |
|  | 10118.3 | - | - | - | 1 | - |
|  | 10126.6 | - | - | - | 1 | - |
|  | 10141.9 | - | - | - | 1 | - |
|  | 10226.2 | - | - | - | 1 | - |
|  | 10296.2 | - | - | - | 1 | - |
|  | 10344.0 | - | - | - | 1 | - |
|  | 10388.4 | - | - | - | 1 | - |
|  | 10430.1 | - | - | - | 1 | - |
|  | 10440.8 | - | - | - | 1 | - |
|  | 13770.4 | - | - | 1 | 1 | 1 |
|  | 13777.6 | - | - | 1 | 1 | 1 |
|  | 13786.8 | - | - | 1 | 1 | 1 |
|  | 13805.6 | - | - | 1 | 1 | 1 |
|  | 13851.0 | - | - | - | 1 | - |
|  | 13853.6 | - | - | - | 1 | - |
|  | 13861.8 | - | - | - | 1 | 1 |
|  | 13870.8 | - | - | - | 1 | - |
|  | 13887.0 | - | - | - | 1 | 1 |
|  | 13888.7 | - | - | - | - | 1 |
|  | 13889.8 | - | - | - | 1 | 1 |
|  | 13891.4 | - | - | - | 1 | 1 |
|  | 13892.9 | - | - | - | 1 | 1 |
|  | 13907.1 | - | - | - | 1 | - |
|  | 13909.0 | - | - | - | 1 | - |
|  | 13910.5 | - | - | - | 1 | - |
|  | 13913.5 | - | - | - | 1 | - |
|  | 13997.2 | - | 1 | 1 | 1 | 1 |
|  | 14008.2 | - | 1 | 1 | 1 | 1 |
|  | 14016.7 | - | - | 1 | 1 | 1 |
|  | 14025.3 | - | - | 1 | 1 | 1 |
|  | 14048.3 | - | - | 1 | 1 | 1 |
|  | 14069.3 | - | - | 1 | - | 1 |
|  | 14869.0 | - | - | - | 1 | - |
|  | 15052.6 | - | - | - | 1 | - |
|  | 16879.2 | - | - | - | 1 | - |
| **Week 2** | *m/z* | 1 | 0 | 5 | 1 | 2 |
|  | 4065.3 | 1 | - | - | - | - |
|  | 6479.9 | - | - | 1 | - | 1 |
|  | 6600.4 | - | - | 1 | - | - |
|  | 10951.4 | - | - | - | 1 | - |
|  | 11774.8 | - | - | - | - | 1 |
|  | 12975.8 | - | - | 1 | - | - |
|  | 13201.2 | - | - | 1 | - | - |
|  | 13204.2 | - | - | 1 | - | - |
| **Week 3** | *m/z* | 0 | 3 | 2 | 2 | 4 |
|  | 5147.2 | - | - | 1 | - | - |
|  | 5436.7 | - | - | - | - | 1 |
|  | 5477.6 | - | - | - | - | 1 |
|  | 7713.0 | - | - | 1 | - | - |
|  | 8179.0 | - | 1 | - | - | - |
|  | 8183.9 | - | 1 | - | - | - |
|  | 8261.0 | - | 1 | - | - | - |
|  | 9962.3 | - | - | - | 1 | 1 |
|  | 10951.4 | - | - | - | 1 | - |
|  | 15052.6 | - | - | - | - | 1 |
| **Week 4** | *m/z* | 0 | 0 | 15 | 0 | 5 |
|  | 5147.2 | - | - | 1 | - | - |
|  | 5676.2 | - | - | 1 | - | - |
|  | 6488.6 | - | - | - | - | 1 |
|  | 6591.4 | - | - | - | - | 1 |
|  | 11296.9 | - | - | 1 | - | - |
|  | 11308.0 | - | - | 1 | - | - |
|  | 11334.3 | - | - | 1 | - | - |
|  | 11349.8 | - | - | 1 | - | - |
|  | 11384.4 | - | - | 1 | - | - |
|  | 12278.0 | - | - | - | - | 1 |
|  | 12284.1 | - | - | - | - | 1 |
|  | 13181.8 | - | - | - | - | 1 |
|  | 15302.8 | - | - | 1 | - | - |
|  | 15309.2 | - | - | 1 | - | - |
|  | 15319.2 | - | - | 1 | - | - |
|  | 15329.3 | - | - | 1 | - | - |
|  | 15330.8 | - | - | 1 | - | - |
|  | 15334.7 | - | - | 1 | - | - |
|  | 15346.5 | - | - | 1 | - | - |
|  | 15359.2 | - | - | 1 | - | - |
| **Week 5** | *m/z* | 0 | 0 | 0 | 0 | 1 |
|  | 8179.0 | - | - | - | - | 1 |
| **Week 6** | *m/z* | 0 | 0 | 0 | 0 | 1 |
|  | 6581.4 | - | - | - | - | 1 |
